# A modification-specific peptide-based immunization approach using CRM197 carrier protein: *Development of a selective vaccine against pyroglutamate Aβ peptides*

**DOI:** 10.1101/084913

**Authors:** Valérie Vingtdeux, Haitian Zhao, Pallavi Chandakkar, Christopher M. Acker, Peter Davies, Philippe Marambaud

**Author notes:** Present address: INSERM, UMR1172 Jean-Pierre Aubert research centre; Université de Lille, Faculté de Médecine; CHRU-Lille, 59045 Lille, France. Corresponding author: Philippe Marambaud.

## Abstract

Strategies aimed at reducing cerebral accumulation of the amyloid-β (Aβ) peptides have therapeutic potential in Alzheimer’s disease (AD). Aβ immunization has proven to be effective at promoting Aβ clearance in animal models but adverse effects have hampered its clinical evaluation. The first anti-Aβ immunization clinical trial, which assessed a full-length Aβ1-42 vaccine, increased the risk of encephalitis most likely because of autoimmune pro-inflammatory T helper 1 (Th1) response against all forms of Aβ. Immunization against less abundant but potentially more pathologically relevant Aβ products, such as N-terminally-truncated pyroglutamate-3 Aβ (AβpE3), could provide efficacy and improve tolerability in Aβ immunotherapy. Here, we describe a selective vaccine against AβpE3 using the diphtheria toxin mutant CRM197 as carrier protein for epitope presentation. CRM197 is currently used in licensed vaccines and has demonstrated excellent immunogenicity and safety in humans. In mice, our AβpE3:CRM197 vaccine triggered the production of specific anti-AβpE3 antibodies that did not cross-react with Aβ1-42, non-cyclized AβE3, or N-terminally-truncated pyroglutamate-11 Aβ (AβpE11). AβpE3:CRM197 antiserum strongly labeled AβpE3 in insoluble protein extracts and decorated cortical amyloid plaques in human AD brains. Anti-AβpE3 antibodies were almost exclusively of the IgG1 isotype, suggesting an anti-inflammatory Th2 response bias to the AβpE3:CRM197 vaccine. To the best of our knowledge, this study shows for the first time that CRM197 has potential as a safe and suitable vaccine carrier for active and selective immunization against specific protein sequence modifications or conformations, such as AβpE3.

## INTRODUCTION

Anti-amyloid-β (Aβ) immunotherapy is under intense investigation in Alzheimer’s disease (AD) (Lemere 2013; Wisniewski & Goñi 2015). Aβ is the core component of the amyloid plaques, a hallmark of the AD brain, and mutations in its precursor APP or in presenilins—the catalytic components of the Aβ-producing enzyme γ-secretase—cause familial AD (Checler 1995; Marambaud & Robakis 2005; Selkoe 2001). Thus, strategies aimed at preventing or lowering Aβ cerebral accumulation might interfere with AD pathogenesis (Citron 2010). Aβ immunization has proven to be very effective at promoting Aβ clearance, at least in animal models. Preclinical and clinical studies, however, have been hampered by unforeseen side effects. The first clinical trial AN-1792, which evaluated an Aβ1-42/QS21 vaccine, was halted in phase II when about 6% of the patients developed meningoencephalitis (Holmes et al. 2008). Although the exact mechanism that led to acute brain inflammation in this clinical trial remains unclear, it is believed that encephalitis arose from an autoimmune reaction triggered by a vaccine directed against the abundant self-protein Aβ coupled to the strong adjuvant QS21, thus favoring pro-inflammatory T helper 1 (Th1) immune responses (Tabira 2010). In this context, second generation anti-Aβ vaccines were designed to prevent T-cell responses during anti-Aβ immunization. These vaccines tested, for instance, N-terminal epitopes within the Aβ sequence and adjuvants that minimize T-cell engagement and favor B-cell responses (Wisniewski & Goñi 2015). Another approach is passive immunization, which has the advantage to bypass T-cell engagement and allow a better control of monoclonal antibody (mAb) dosage and epitope targeting. However, recent phase III AD trials of two anti-Aβ mAb, solanezumab and bapineuzumab, failed to slow cognitive or functional decline in patients with mild-to-moderate AD (Hampel et al. 2015). The main argument put forward to explain the lack of efficacy of these passive immunization approaches was that treatment might have started too late to reverse or delay the disease process (Karran & Hardy 2014). It is also possible that passive immunization might not deliver enough mAb to promote plaque clearance. Passive immunization also raised concerned because of the practical and financial sustainability of injecting and monitoring mAb injections on a regular basis for several years (Golde 2014).

In the amyloidogenic pathway, APP is sequentially endoproteolyzed by the proteases β-secretase/BACE1 and the presenilin/γ-secretase complex to produce various Aβ peptides, including the most abundant isoforms Aβ1-40 and Aβ1-42 (De Strooper et al. 2010). In addition to these major Aβ isoforms, N-terminally truncated Aβ products have been identified in the AD brain, including peptides starting with pyroglutamate residues at the position 3 (AβpE3) and 11 (AβpE11) (Mori et al. 1992; Portelius et al. 2010; Saido et al. 1995). N-terminal truncation was proposed to be mediated, at least in part, by aminopeptidase A (Sevalle et al. 2009) and cyclization of N-terminally-exposed glutamates is catalyzed by the enzyme glutaminyl cyclase (Schilling et al. 2008). Pyroglutamate Aβ is a promising target because appears to play a key role in Aβ oligomerization, seeding, and stabilization (Dammers et al. 2015; He & Barrow 1999; Nussbaum et al. 2012; Piccini et al. 2005; Schilling et al. 2006; Schlenzig et al. 2009). Furthermore, pyroglutamate Aβ has specific neurotoxic properties in cell cultures and leads to cerebral neuronal loss and synaptic function impairments in mice (Alexandru et al. 2011; Gunn et al. 2016; Russo et al. 2002; Schlenzig et al. 2012).

In this context, recent studies have proposed that immunization against less abundant but potentially more amyloidogenic and more neurotoxic isoforms of Aβ, such as AβpE3, could improve tolerability and efficacy in Aβ immunotherapy (Bayer & Wirths 2014; Cynis et al. 2016; Frost et al. 2015). These studies, however, are so far limited to passive immunization using antibodies previously screened *in vitro* for their specificity to AβpE3 and non-cross-reactivity to full length Aβ. Here, we describe a novel AβpE3 vaccine using the non-toxic mutant of diphtheria toxin CRM197 (cross-reacting material 197) as carrier protein for epitope presentation. CRM197 has been extensively used in licensed vaccines directed against capsular polysaccharides of several bacterial pathogens and has demonstrated excellent efficacy and tolerability in humans (Bröker et al. 2011; Shinefield 2010). Because recent data have shown that CRM197 is also suitable for conjugation to, and presentation of, peptides (Caro-Aguilar et al. 2013), we speculated that CRM197 could be conjugated to minimal peptide epitopes to facilitate the generation of conformation/modification-specific antibodies directed against pyroglutamate Aβ. We show that our vaccines in mice—composed of AβpE3-8 or AβpE11-16 peptides covalently conjugated to CRM197—triggered the production of fully specific antibodies directed against AβpE3 and AβpE11, respectively. Anti-AβpE3 antibodies stained brains from AD patients by western blotting (WB) and immunohistochemistry (IHC), and were almost exclusively of the IgG1 isotype, indicating the engagement of a Th2 response.

## MATERIALS AND METHODS

### Preparation of the AβpE3-8:CRM197 and AβpE11-16:CRM197 conjugate vaccines

N-terminally acetylated and C-terminally amidated AβpE3-8 and AβpE11-16 peptides containing a C-terminal cysteine residue preceded by a two-glycine-bridge (pEFRHDSGGC and pEVHHQKGGC, respectively; GenScript) were solubilized in phosphate buffered saline (PBS) containing 2 mM EDTA (PBS/EDTA) to obtain 4 mg/mL solutions. CRM197 (List Biological Labs) was reconstituted with PBS/EDTA to obtain a 2 mg/mL solution. To irreversibly crosslink the peptides to the carrier protein, CRM197 was first activated with a 20-fold molar excess of succinimidyl-4-(N-maleimidomethyl)cyclohexane-1-carboxylate (SMCC) crosslinker (Pierce, at 1.5 mg/mL in DMSO). After a 30 min incubation at room temperature (RT), the SMCC/CRM197 mixture was desalted on a resin column (Zeba Desalt Spin Column, Pierce). Activated CRM197 was then combined with AβpE3-8 and AβpE11-16 peptide solutions and incubated for 30 min at RT. A ratio of 1:10 (CRM197 to peptides) was empirically chosen. Successful conjugation was confirmed by SDS-PAGE and coomassie staining (GelCode, Pierce, see Fig. 2A).

### Immunization and sample collection

Animal experiments were performed according to procedures approved by the Feinstein Institute for Medical Research Institutional Animal Care and Use Committee. C57BL/6J male mice (~3-month-old) were injected subcutaneously with 10 μg of AβpE3-8:CRM197 and AβpE11-16:CRM197 conjugate vaccines or with 10 μg of free AβpE3-8 and AβpE11-16 peptides, used as negative controls. A prime injection (Day 0) was followed by a boost injection at Day 14 of the same amount of vaccines or free peptides. Blood samples were taken at Day 0 and Day 30.

### ELISA for measurement of antibody responses

Anti-AβpE3-8, anti-AβpE11-16, and anti-CRM197 antibody levels in mouse serum were determined by enzyme-linked immunosorbent assay (ELISA). 96-well plates (Maxisorp, Nunc) were coated with 100 μL of 2 μg/mL AβpE3-8, AβpE11-16, or CRM197 in carbonate buffer (0.05 M, pH 9.6) and incubated overnight at 4^o^C. The following morning, plates were blocked for 1 h at RT with 5% skim milk in TBS containing 0.05% Tween 20 (TBST). After washing with TBST, serial dilutions of individual mouse serum samples (diluted in PBST containing 1% skim milk) were prepared and 100 μL/well of mouse serum was incubated for 2 h at RT. After 5 more washes, 100 μL/well horseradish peroxidase (HRP)-conjugated goat anti-mouse immunoglobins (Igs) secondary antibody (Southern Biotech, diluted 1:500 in PBST containing 1% skim milk) was incubated for 1 h at RT. TMB substrate was added after another wash and the reaction was allowed to develop for 30 min at RT. The optical density was measured at 405 nm using a TECAN GENios Pro plate reader. Antibody responses were expressed as titers. Antibody titers were determined via a linear fit for optical density values of 9 dilutions, and expressed in arbitrary units by calculating the reciprocal dilution that gave 50% of the maximum absorbance response. For measurement of Ig isotype specificity (IgG1, IgG2b, IgG2c, IgG3, IgA, and IgM), an ELISA protocol similar to that described above was followed. Briefly, ELISA plates were coated as described above. Reference mouse IgG1, IgG2b, IgG2c, IgG3, IgA and IgM antibodies were diluted in TBST containing 1% skim milk. To detect specific binding, HRP-conjugated anti-mouse IgG1, IgG2b, IgG2c, IgG3, IgA and IgM antibodies (Southern Biotech) were used at a dilution of 1:500, TMB was then added as described above. The presence of antigen-specific isotype-specific antibodies was detected and measured as described above. Antibody levels were expressed in micrograms per milliliter.

### Statistical analysis

For comparison of results within experimental groups, a Student’s t test was performed. For all multigroup comparisons, one-way analysis of variance (ANOVA) and post-hoc Bonferoni test for multiple comparisons were used. Statistical significance was defined as p < 0.05.

### Human cases and brain protein extraction

Cases were obtained from the Albert Einstein College of Medicine human brain bank, Bronx, NY (see Table). Brain samples (see Table) were processed as previously described (Vingtdeux et al. 2011). Briefly, human brains were sequentially extracted to obtain a soluble SDS fraction and an insoluble formic acid (FA) fraction. Samples were first homogenized and sonicated in Tris-buffered saline (TBS) containing 2% SDS and 1x Complete protease inhibitor mixture (Roche Applied Science) and centrifuged at 100,000 x *g* for 1 h at 4°C. The supernatant was removed and the resulting pellet was then extracted with 70% formic acid in water. FA extracts were dried under vacuum in a speed vacuum and thus dissolved in DMSO.

### Western blot (WB)

Protein extracts from SDS and FA fractions obtained from human brain samples were analyzed by SDS-PAGE using the indicated antibodies. Samples were electrophoresed on 10% Tris-HCl gels (phospho-tau and actin analysis) or on 16.5% Tris-Tricine gels (Aβ isoforms, Biorad) and transferred onto nitrocellulose membranes. Aβ was detected as previously described (Vingtdeux et al. 2015). Briefly, membranes were microwaved for 5 min in PBS. Membranes were then blocked in 5% fat-free milk in TBS, and incubated with primary antibodies overnight at 4°C. A standard ECL detection procedure was then used.

### Immunohistochemistry (IHC)

Five-μm-thick sections of formalin-fixed paraffin-embedded brain tissue were immunostained with 6E10 mAb (Covance, 1:1000 dilution) or AβpE3:CRM197 antiserum (1:2 dilution). Sections were deparaffinized by immersion in xylene and hydration through graded ethanol solutions. Denaturation for antigen recovery was performed by incubation of the slides in 70% formic acid for 30 min at RT. Denaturation was stopped by incubation in 100 mM Tris-HCl, pH 7.4 for 5 min. After washing once in TBS containing 0.05% Triton-X100 (TBSTx), endogenous peroxidase activity was inhibited by incubation in 5% hydrogen peroxide in TBSTx for 30 min at RT. After washing twice in TBSTx for 5 min and once in water, sections were blocked in 10% fat-free milk (6E10) or 5% normal goat serum (AβpE3:CRM197 antiserum) in TBSTx containing 1 mg/mL BSA and 1 mM NaF for 1 h at RT. Sections were then incubated in the presence of primary antibodies diluted in 10% fat-free milk (6E10) or 5% normal goat serum (AβpE3:CRM197 antiserum) in TBSTx containing 1 mg/ml BSA and 1 mM NaF overnight at 4^o^C in a humidified chamber. After washing, the sections were incubated with biotin-coupled anti-mouse IgG1 secondary antibodies (1:1,000 dilution in TBSTx with 20% Superblock, Pierce) before incubation with streptavidin-HRP (1:1,000 dilution in TBSTx with 20% Superblock, Southern Biotech) and visualization with diaminobenzidine tetrahydrochloride.

## RESULTS

### Immunogenicity of CRM197 in mice

We first verified that unconjugated CRM197 without adjuvant was immunogenic in C57BL/6J mice. Based on previously published dose-ranging immunogenicity analyses of different CRM197-based vaccines (Caro-Aguilar et al. 2013; Rondini et al. 2011), we choose to inject 10 μg of CRM197. A prime injection of unconjugated CRM197 followed by a boost injection at Day 15 of the same amount of carrier protein elicited robust immunogenicity against CRM197 at Day 30 (Fig. 1A). Saline control injections as expected did not elicit an anti-CRM197 antibody response (Fig. 1A). Ig isotype determination revealed that anti-CRM197 antibodies were exclusively of the IgG1 isotype and reached levels of ~35 μg/mL at Day 30 post-prime injection (Fig. 1B).

**Figure 1:**
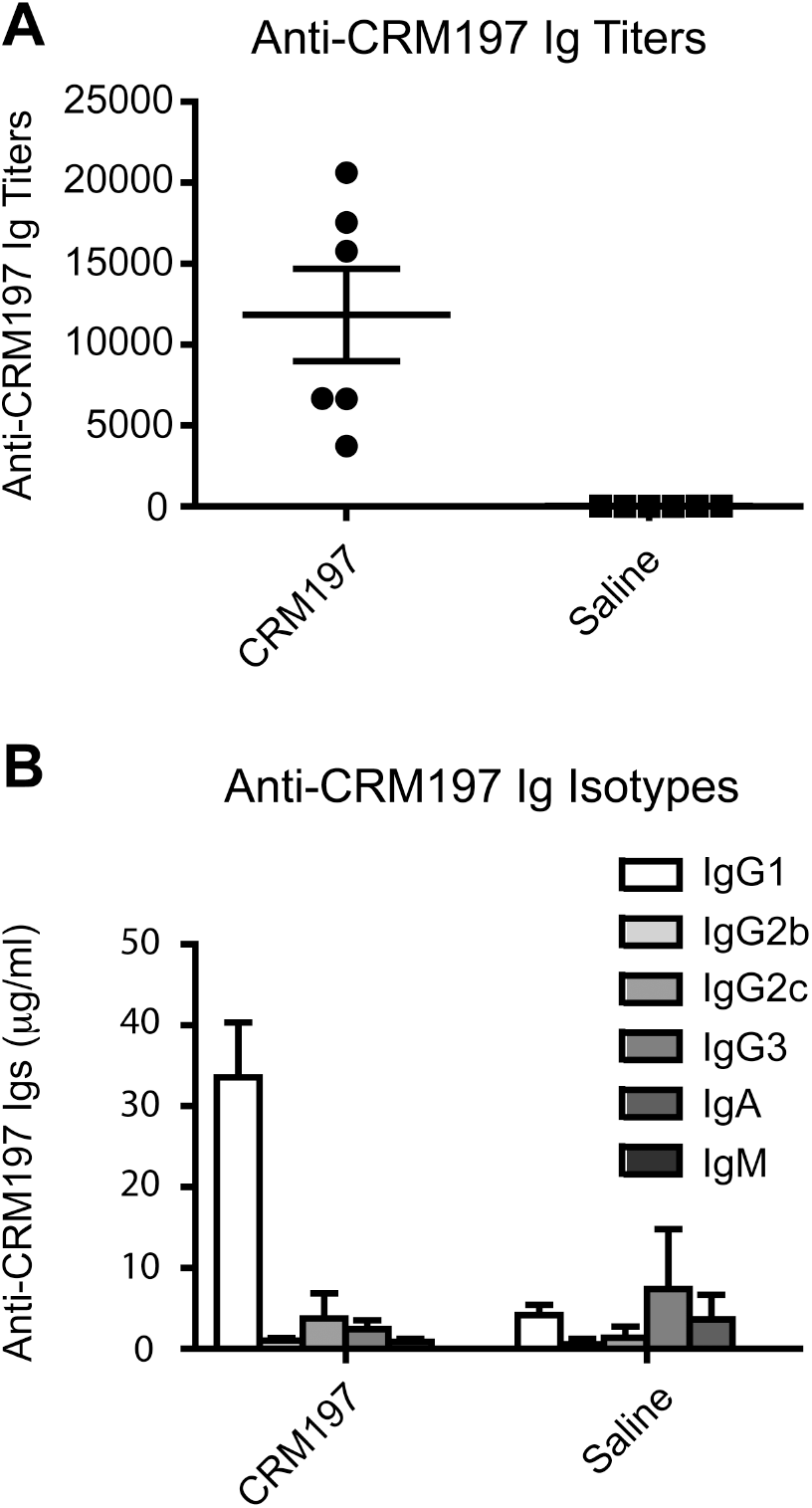
Anti-CRM197 antibody response in mice immunized with CRM197. Specific anti-CRM197 Ig titers (A) and Ig isotype concentration (IgG1, IgG2a, IgG2b, IgG3, IgA, IgM) (B) quantified by ELISA in sera obtained from C57BL/6J mice immunized (CRM197) or not (Saline) with CRM197. Sera were collected 30 days after prime injection. Data are mean ± SEM from 6 mice per group, each dot represent individual mouse serum.

### Immunogenicity of the AβpE3:CRM197 and AβpE11:CRM197 conjugate vaccines

A short peptide sequence of 6 residues starting at pE3 (pE3-8) or pE11 (pE11-16) linked in C-terminal to a 3-residue spacer (see Methods) was chosen to facilitate the production of conformation/modification-specific antibodies directed against Aβ. A 6-residue peptide epitope has also the advantage to be shorter than conventional T-cell epitopes and thus might prevent unwanted T-cell activation and detrimental pro-inflammatory responses (Rammensee 1995). Molecular weight analysis by protein staining after SDS-PAGE revealed that pE3-8 and pE11-16 peptides could be covalently conjugated to CRM197 (Fig. 2A). The generated AβpE3:CRM197 and AβpE11:CRM197 conjugates (10 μg) elicited robust antibody responses against AβpE3-8 and AβpE11-16 peptides, respectively, when assessed by ELISA. Total serum Ig titers were in average of 1,000 at Day 30 after one prime injection and one boost injection at Day 15 (Figs. 2B and 2D). Importantly, the vaccines did not generate antibodies directed against the corresponding non-cyclized Aβ3-8 and Aβ11-16 peptides (Figs. 2C and 2E), showing that immunogenicity was specific to the pE modification. No anti-AβpE3- or anti-AβpE11-specific antibodies were detected in control mice injected with unconjugated CRM197 or saline (Figs. 2B and 2D). Injection of free AβpE3-8 and AβpE11-16 peptides (unconjugated to CRM197) did not elicit immunogenic responses against these peptides (Figs. 2B and 2D).

**Figure 2:**
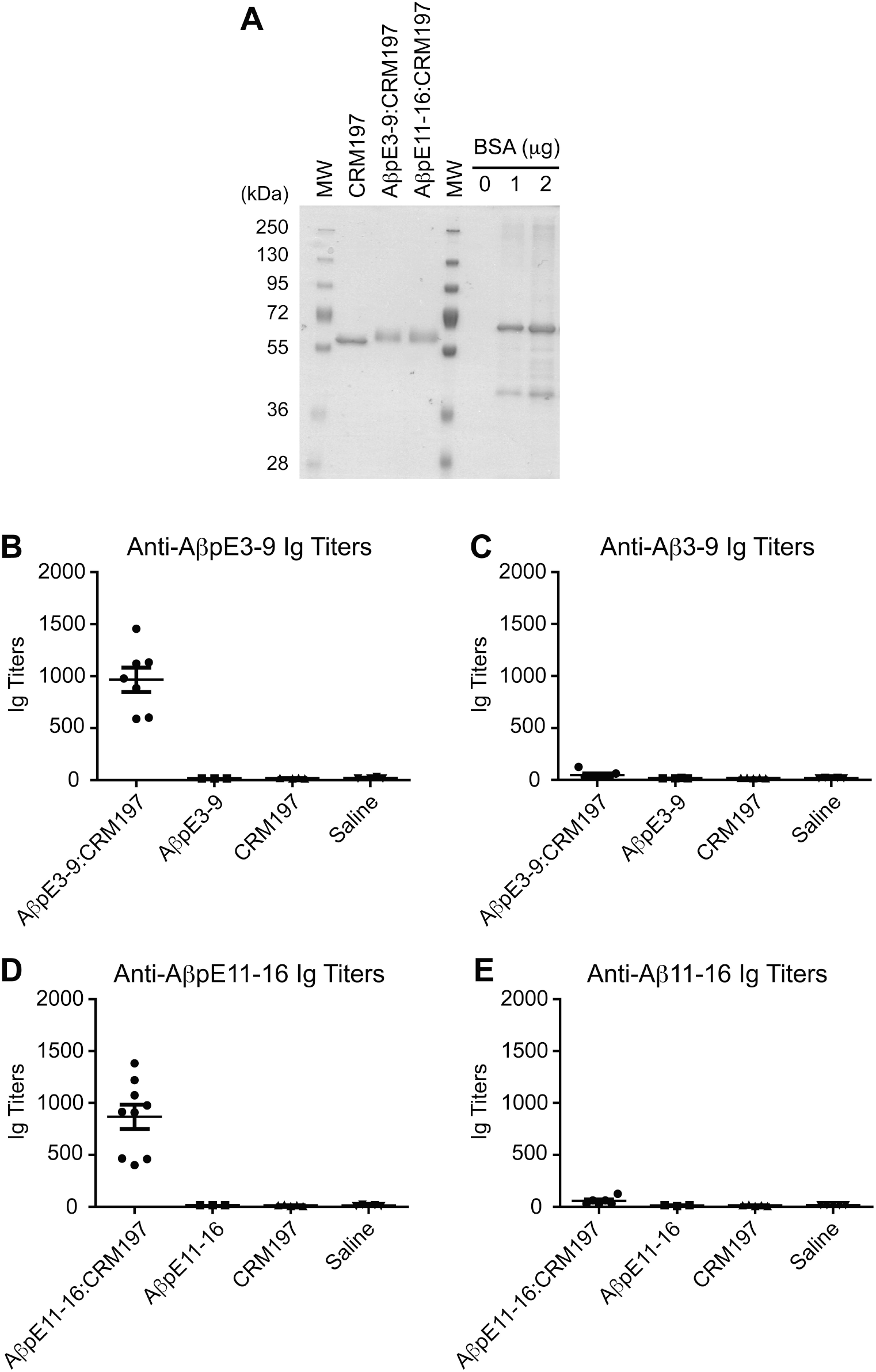
Anti-AβpE3-9 and anti-AβpE11-16 antibody response. Coomassie blue staining of CRM197 alone or conjugated to AβpE3-9 (AβpE3-9:CRM197) or AβpE11-16 (AβpE11-16:CRM197) (A). Anti-AβpE3-9 (B), anti-Aβ3-9 (C), anti-AβpE11-16 (D), and anti-Aβ11-16 (E) Ig titers in sera obtained from mice immunized with AβpE3-9 or AβpE11-16 conjugated (AβpE3-9:CRM197, AβpE11-16:CRM197) or not (AβpE3-9, AβpE11-16) with CRM197. Data represents mean ± SEM from 3-9 mice per group. MW, molecular weight makers.

As observed for the immunogenic responses to unconjugated CRM197 (Fig. 1), the anti-AβpE3 and anti-AβpE11 antibodies were exclusively of the IgG1 isotype (Figs. 3A and 3B). At Day 30 post-prime injection, AβpE3:CRM197 and AβpE11:CRM197 vaccines elicited the production of ~1.5 μg/mL of serum specific IgG1.

**Figure 3:**
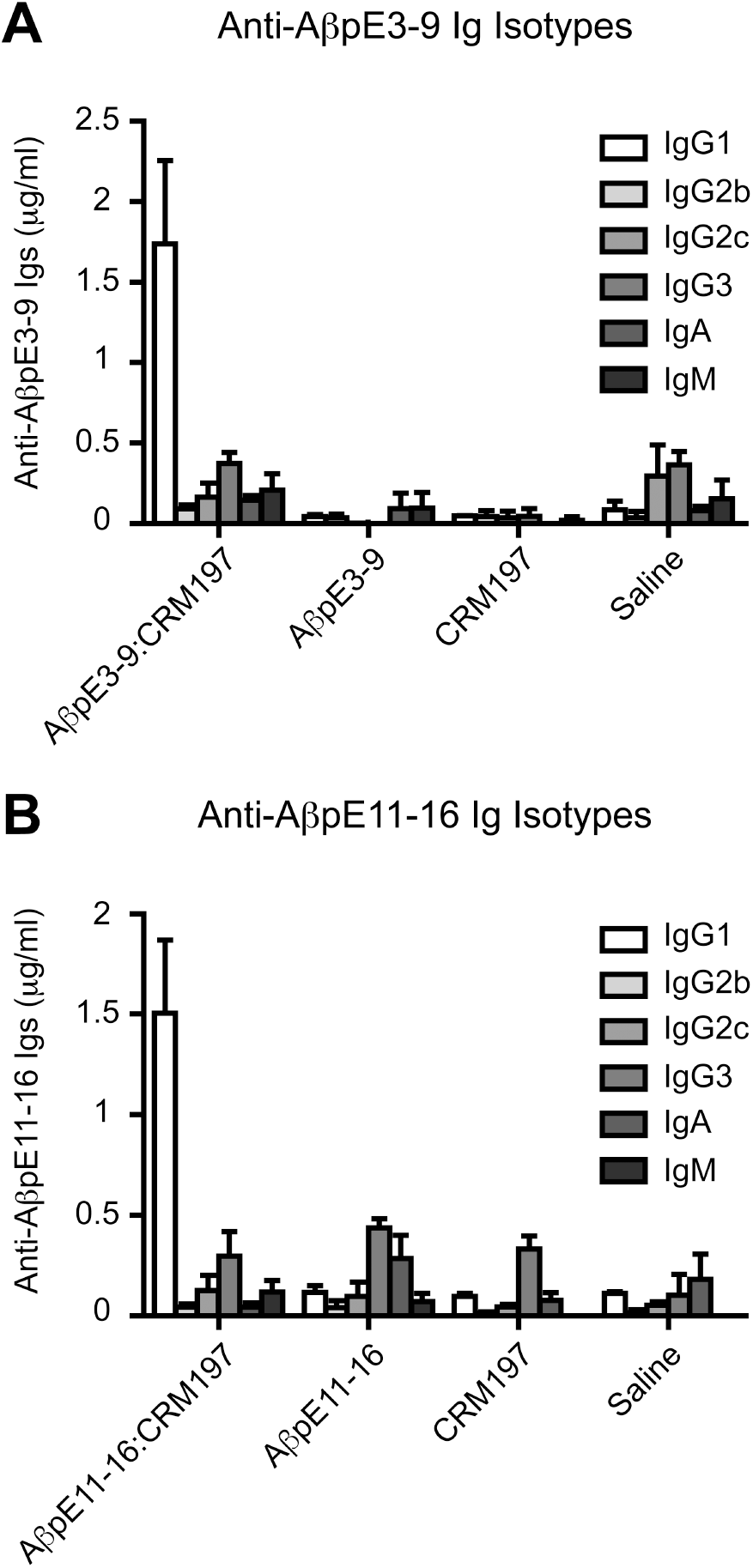
Isotype specificity of the Ig response elicited by AβpE3-9:CRM197 and AβpE11-16:CRM197 vaccines. Anti-AβpE3-9 (A) or anti-AβpE11-16 Ig isotype (IgG1, IgG2a, IgG2b, IgG3, IgA, IgM) concentrations in sera obtained from mice immunized with AβpE3-9 or AβpE11-16 conjugated (AβpE3-9:CRM197, AβpE11-16:CRM197) or not (AbpE3-9, AbpE11-16) with CRM197 (B). Data represents mean ± SEM from 3-7 mice per group.

### The AβpE3:CRM197 vaccine produces antibodies that react with amyloid deposits in AD brains

Antiserum produced after immunization with the AβpE3:CRM197 vaccine labeled synthetic AβpE3-42 peptide by WB (Fig. 4A). Importantly, AβpE3:CRM197 antiserum did not cross-react with synthetic AβpE11-42 or synthetic Aβ1-42 (Fig. 4A), showing that the antibodies produced did not recognize non-specifically the pE modification nor they interacted with an internal epitope containing residues 3-8 of Aβ1-42. AβpE3:CRM197 antiserum is thus fully specific to 241 AβpE3. Strikingly, AβpE3:CRM197 antiserum was also able to label AβpE3 in insoluble/aggregated brain protein preparations (FA fractions, see Methods) obtained from 6 independent well-characterized AD patients (see Table). Almost no immunoreactivity was observed in the corresponding soluble preparations (SDS fractions) (Fig. 4B), indicating that the anti-AβpE3 antibodies produced by the AβpE3:CRM197 vaccine recognize human amyloid 246 material in the AD brain, and that AβpE3 is mostly aggregated. Of note, AβpE3 immunoreactivity was also observed in normal control brains that contained large amounts of aggregated total Aβ (also detected with 6E10 mAb in the insoluble fractions) (Fig. 4B). In fact, a strong parallel was observed between the levels of aggregated total Aβ and aggregated AβpE3 in brains from both normal and AD individuals (Fig. 4B), suggesting that AβpE3 is a marker for amyloid deposition but is not specific to AD.

**Figure 4:**
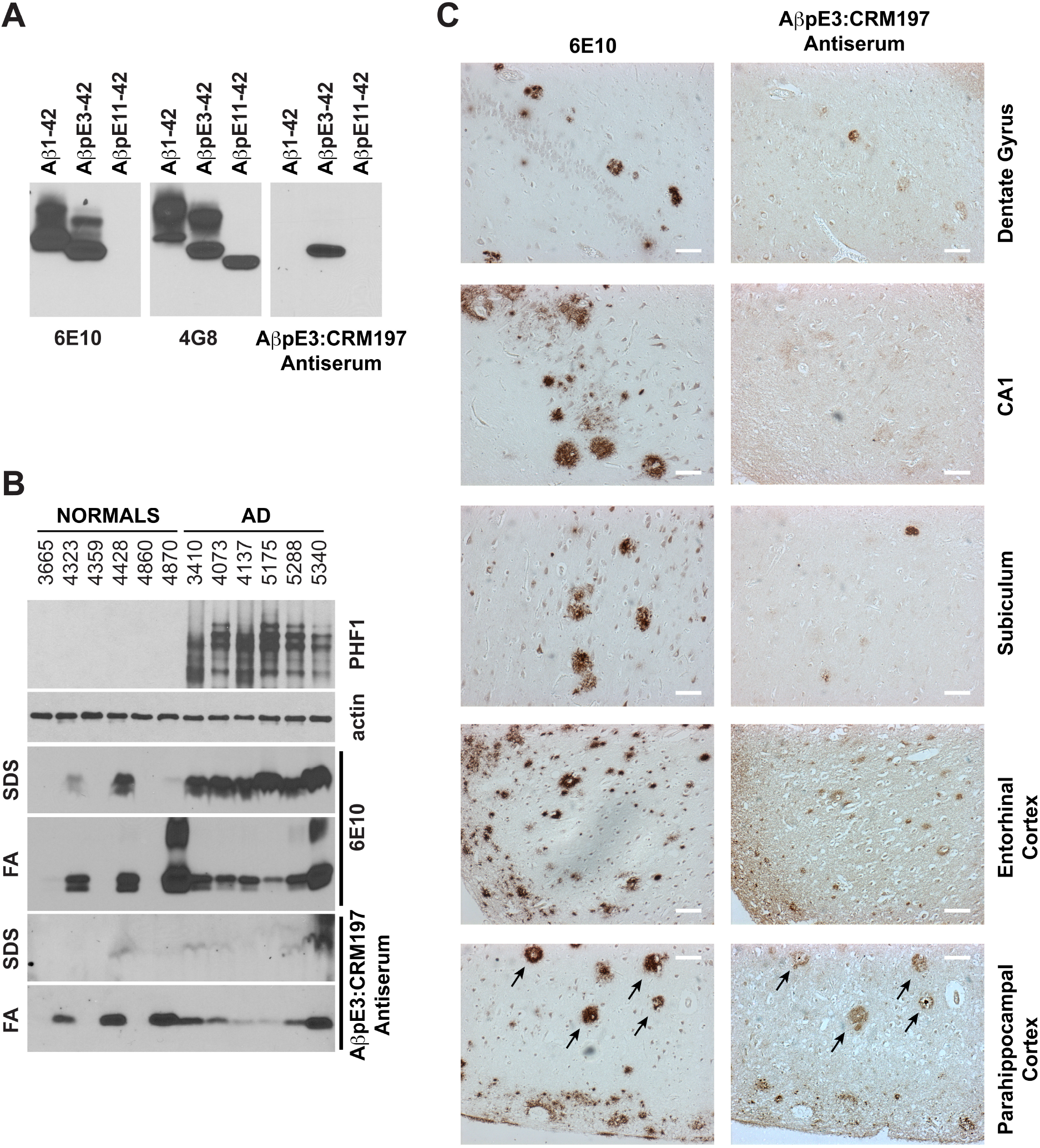
AβpE3-9:CRM197 antiserum characterization. WB analysis of recombinant Aβ1-42, AβpE3-42, and AβpE11-42 using 6E10, 4G8, or AβpE3-9:CRM197 antiserum showing the specificity of the AβpE3-9:CRM197 antiserum towards AβpE3-42 (A). WB analysis of human brain homogenates obtained from normal or AD cases (Braak stage V-VI, see Table) using PHF1, actin, and 6E10 antibodies, or AβpE3-9:CRM197 antiserum (B). IHC of serial brain sections from an AD case stained with 6E10 antibody or AβpE3-9:CRM197 antiserum (C).

In addition, AβpE3:CRM197 antiserum decorated amyloid plaques in human AD brain slices by IHC. Most of the anti-AβpE3 immunoreactivity was observed on plaques present in the cortex (entorhinal and parahippocampal cortices, Fig. 4C). Although amyloid plaque deposition was very pronounced in the AD case analyzed, only a few plaques were labeled with the AβpE3:CRM197 antiserum in the hippocampal formation (the dentate gyrus, CA1, and subiculum, Fig.4C). Interestingly, in adjacent sections of the parahippocampal cortex, identical plaques stained for both total Aβ (6E10 mAb) and AβpE3 (Fig. 4C, arrows). This co-staining revealed that anti-AβpE3 antiserum preferentially decorated highly dense plaques. Of note, anti-IgG1 secondary antibodies were used in combination with AβpE3:CRM197 antiserum for WB and IHC further showing that the immunoreactive anti-AβpE3 antibodies are of the IgG1 isotype.

## DISCUSSION

In this study, we show that minimal epitopes can be designed to generate vaccines that are fully specific to the pyroglutamate modifications of Aβ. Indeed, antiserum obtained after immunization with the AβpE3:CRM197 conjugate vaccine, demonstrated excellent immunoreactivity against synthetic and AD brain-derived AβpE3, with no detectable cross-reactivity with Aβ1-42, non-cyclized AβE3, or AβpE11. Further analysis using AD brain samples revealed that AβpE3:CRM197 antiserum mainly decorated highly aggregated amyloid material, supporting the notion that AβpE3 is associated with the dense core of the senile plaques (Sullivan et al. 2011).

CRM197 is routinely used as conjugate vaccine in licensed vaccines directed against bacterial capsular polysaccharides (Bröker et al. 2011; Shinefield 2010). The use of CRM197 for peptide epitope presentation has not been approved yet for human use, but pre-clinical evidence has already suggested that it has therapeutic potential (Caro-Aguilar et al. 2013). In addition, an experimental vaccine against Aβ using CRM197 is currently being evaluated in a Phase II trial (ACC-001, Elan/J&J/Wyeth). In the ACC-001 vaccine, CRM197 is conjugated to the N-terminal residues 1-6 of Aβ and thus is aimed at targeting all Aβ isoforms containing this N-end of Aβ (Winblad et al. 2014). Our study strengthens the notion that CRM197 has strong potential for peptide presentation. This work further demonstrates that CRM197 might particularly be useful for the generation of conformation/modification-specific anti-peptide vaccines.

Both active and passive immunization directed against Aβ have their respective pros and cons (see Introduction) and there is no consensus yet as to whether one approach has a stronger therapeutic potential for AD. Both approaches—targeting different Aβ epitopes—are therefore actively investigated at the pre-clinical and clinical levels. Our approach was aimed at increasing the anti-Aβ immunotherapy toolkit by proposing an active immunization strategy specifically targeting N-terminally-truncated pyroglutamate Aβ. Further studies will be required to validate the utility of such a vaccine in AD mouse models of amyloid deposition. It should be noted, however, that several studies using passive immunization specifically targeting AβpE3 have already been conducted in different mouse amyloid models. These studies provided solid and concordant evidence that anti-AβpE3 antibodies have an overall plaque lowering effect and can improve the associated cognitive/behavioral deficits (Demattos et al. 2012; Frost et al. 2015; Wirths et al. 2010). Thus, it will be important to determine whether selective and active anti-AβpE3 immunization could also reduce the pathology in these models.

In this context, new immunization formulations for the AβpE3:CRM197 vaccine will have to be tested to boost antibody production. Indeed, although the AβpE3:CRM197 antiserum was able to react with both synthetic AβpE3 and AD-brain derived amyloid material, its antibody levels in the serum around 1.5 μg/mL are likely to be too low to allow clearance of AβpE3 and amyloid in mouse AD models. Thus, future experiments will also have to determine whether immunogenicity of the AβpE3:CRM197 vaccine can be improved by repeated injections and/or addition of adjuvants. Interestingly, recent data demonstrated that combined addition of the adjuvants aluminum hydroxide and CpG can increase more than 100-fold the antibody titer of a nicotine:CRM197 conjugate vaccine (McCluskie et al. 2015) and QS-21 elicited consistently higher anti-Aβ IgG titers in a phase IIa trial for the Aβ1-6:CRM197 ACC-001 vaccine (Pasquier et al. 2016).

In conclusion, we propose that conjugation of peptides AβpE3-8 and AβpE11-16 to CRM197 can generate fully specific vaccines directed against AβpE3 and AβpE11, respectively. The main strength of this immunization strategy is to combine the advantages of a vaccine with the high specificity of targeting particularly amyloidogenic and neurotoxic sub-species of Aβ, while sparing the more abundant and maybe more physiologically relevant full-length Aβ40 and Aβ42.

---

**Table:** Human cases analyzed in this study. AD, Alzheimer’s disease; Mid-Temp Cortex, mid-temporal cortex; Hp, hippocampal region. Cases were obtained from the Albert Einstein College of Medicine human brain bank, Bronx, NY.

| Case # | Clinical Dx | Braak | Age | Sex | Region |
| --- | --- | --- | --- | --- | --- |
| 3665 | Normal | 0 | 85 | Female | Mid-Temp Cortex |
| 4323 | Normal | 0 | 93 | Female | Mid-Temp Cortex |
| 4359 | Normal | 0 | 92 | Male | Mid-Temp Cortex |
| 4428 | Normal | 0 | 84 | Female | Mid-Temp Cortex |
| 4860 | Normal | 0 | 92 | Female | Mid-Temp Cortex |
| 4870 | Normal | 0 | 99 | Female | Mid-Temp Cortex |
| 3410 | AD | V-VI | 81 | Male | Mid-Temp Cortex |
| 4073 | AD | V-VI | 78 | Female | Mid-Temp Cortex |
| 4137 | AD | V-VI | 88 | Male | Mid-Temp Cortex |
| 5175 | AD | V-VI | 56 | Female | Mid-Temp Cortex |
| 5288 | AD | V-VI | 81 | Male | Mid-Temp Cortex |
| 5340 | AD | V-VI | 54 | Male | Mid-Temp Cortex |

## REFERENCES

Alexandru A, Jagla W, Graubner S, Becker A, Bäuscher C, Kohlmann S, Sedlmeier R, Raber KA, Cynis H, Rönicke R, Reymann KG, Petrasch-Parwez E, Hartlage-Rübsamen M, Waniek A, Rossner S, Schilling S, Osmand AP, Demuth HU, and von Hörsten S. 2011. Selective hippocampal neurodegeneration in transgenic mice expressing small amounts of truncated Aβ is induced by pyroglutamate-Aβ formation. J Neurosci 31:12790-12801. 10.1523/JNEUROSCI.1794-11.2011

Bayer TA, and Wirths O. 2014. Focusing the amyloid cascade hypothesis on N-truncated Abeta peptides as drug targets against Alzheimer's disease. Acta Neuropathol 127:787-801. 10.1007/s00401-014-1287-x

Bröker M, Costantino P, DeTora L, McIntosh ED, and Rappuoli R. 2011. Biochemical and biological characteristics of cross-reacting material 197 CRM197, a non-toxic mutant of diphtheria toxin: use as a conjugation protein in vaccines and other potential clinical applications. Biologicals 39:195-204. 10.1016/j.biologicals.2011.05.004

Caro-Aguilar I, Ottinger E, Hepler RW, Nahas DD, Wu C, Good MF, Batzloff M, Joyce JG, Heinrichs JH, and Skinner JM. 2013. Immunogenicity in mice and non-human primates of the Group A Streptococcal J8 peptide vaccine candidate conjugated to CRM197. Hum Vaccin Immunother 9:488-496.

Checler F. 1995. Processing of the beta-amyloid precursor protein and its regulation in Alzheimer's disease. J Neurochem 65:1431-1444.

Citron M. 2010. Alzheimer's disease: strategies for disease modification. Nat Rev Drug Discov 9:387-398. 10.1038/nrd2896

Cynis H, Frost JL, Crehan H, and Lemere CA. 2016. Immunotherapy targeting pyroglutamate-3 Aβ: prospects and challenges. Mol Neurodegener 11:48. 10.1186/s13024-016-0115-2

Dammers C, Gremer L, Reiß K, Klein AN, Neudecker P, Hartmann R, Sun N, Demuth HU, Schwarten M, and Willbold D. 2015. Structural Analysis and Aggregation Propensity of Pyroglutamate Aβ(3-40) in Aqueous Trifluoroethanol. PLoS One 10:e0143647. 10.1371/journal.pone.0143647

De Strooper B, Vassar R, and Golde T. 2010. The secretases: enzymes with therapeutic potential in Alzheimer disease. Nat Rev Neurol 6:99-107. 10.1038/nrneurol.2009.218

Demattos RB, Lu J, Tang Y, Racke MM, Delong CA, Tzaferis JA, Hole JT, Forster BM, McDonnell PC, Liu F, Kinley RD, Jordan WH, and Hutton ML. 2012. A plaque-specific antibody clears existing β-amyloid plaques in Alzheimer's disease mice. Neuron 76:908-920. 10.1016/j.neuron.2012.10.029

Frost JL, Liu B, Rahfeld JU, Kleinschmidt M, O'Nuallain B, Le KX, Lues I, Caldarone BJ, Schilling S, Demuth HU, and Lemere CA. 2015. An anti-pyroglutamate-3 Aβ vaccine reduces plaques and improves cognition in APPswe/PS1ΔE9 mice. Neurobiol Aging 36:3187-3199. 10.1016/j.neurobiolaging.2015.08.021

Golde TE. 2014. Open questions for Alzheimer's disease immunotherapy. Alzheimers Res Ther 6:3. 10.1186/alzrt233

Gunn AP, Wong BX, Johanssen T, Griffith JC, Masters CL, Bush AI, Barnham KJ, Duce JA, and Cherny RA. 2016. Amyloid-β Peptide Aβ3pE-42 Induces Lipid Peroxidation, Membrane Permeabilization, and Calcium Influx in Neurons. J Biol Chem 291:6134-6145. 10.1074/jbc.M115.655183

Hampel H, Schneider LS, Giacobini E, Kivipelto M, Sindi S, Dubois B, Broich K, Nisticò R, Aisen PS, and Lista S. 2015. Advances in the therapy of Alzheimer's disease: targeting amyloid beta and tau and perspectives for the future. Expert Rev Neurother 15:83-105. 10.1586/14737175.2015.995637

He W, and Barrow CJ. 1999. The A beta 3-pyroglutamyl and 11-pyroglutamyl peptides found in senile plaque have greater beta-sheet forming and aggregation propensities in vitro than full-length A beta. Biochemistry 38:10871-10877. 10.1021/bi990563r

Holmes C, Boche D, Wilkinson D, Yadegarfar G, Hopkins V, Bayer A, Jones RW, Bullock R, Love S, Neal JW, Zotova E, and Nicoll JA. 2008. Long-term effects of Abeta42 immunisation in Alzheimer's disease: follow-up of a randomised, placebo-controlled phase I trial. Lancet 372:216-223. 10.1016/S0140-6736(08)61075-2

Karran E, and Hardy J. 2014. A critique of the drug discovery and phase 3 clinical programs targeting the amyloid hypothesis for Alzheimer disease. Ann Neurol 76:185-205. 10.1002/ana.24188

Lemere CA. 2013. Immunotherapy for Alzheimer's disease: hoops and hurdles. Mol Neurodegener 8:36. 10.1186/1750-1326-8-36

Marambaud P, and Robakis NK. 2005. Genetic and molecular aspects of Alzheimer's disease shed light on new mechanisms of transcriptional regulation. Genes Brain Behav 4:134-146. 10.1111/j.1601-183X.2005.00086.x

McCluskie MJ, Thorn J, Gervais DP, Stead DR, Zhang N, Benoit M, Cartier J, Kim IJ, Bhattacharya K, Finneman JI, Merson JR, and Davis HL. 2015. Anti-nicotine vaccines: Comparison of adjuvanted CRM197 and Qb-VLP conjugate formulations for immunogenicity and function in non-human primates. Int Immunopharmacol 29:663-671. 10.1016/j.intimp.2015.09.012

Mori H, Takio K, Ogawara M, and Selkoe DJ. 1992. Mass spectrometry of purified amyloid beta protein in Alzheimer's disease. J Biol Chem 267:17082-17086.

Nussbaum JM, Schilling S, Cynis H, Silva A, Swanson E, Wangsanut T, Tayler K, Wiltgen B, Hatami A, Rönicke R, Reymann K, Hutter-Paier B, Alexandru A, Jagla W, Graubner S, Glabe CG, Demuth HU, and Bloom GS. 2012. Prion-like behaviour and tau-dependent cytotoxicity of pyroglutamylated amyloid-β. Nature 485:651-655. 10.1038/nature11060

Pasquier F, Sadowsky C, Holstein A, Leterme GL, Peng Y, Jackson N, Fox NC, Ketter N, Liu E, and Ryan JM. 2016. Two Phase 2 Multiple Ascending-Dose Studies of Vanutide Cridificar (ACC-001) and QS-21 Adjuvant in Mild-to-Moderate Alzheimer's Disease. J Alzheimers Dis. 10.3233/jad-150376

Piccini A, Russo C, Gliozzi A, Relini A, Vitali A, Borghi R, Giliberto L, Armirotti A, D'Arrigo C, Bachi A, Cattaneo A, Canale C, Torrassa S, Saido TC, Markesbery W, Gambetti P, and Tabaton M. 2005. beta-amyloid is different in normal aging and in Alzheimer disease. J Biol Chem 280:34186-34192. 10.1074/jbc.M501694200

Portelius E, Bogdanovic N, Gustavsson MK, Volkmann I, Brinkmalm G, Zetterberg H, Winblad B, and Blennow K. 2010. Mass spectrometric characterization of brain amyloid beta isoform signatures in familial and sporadic Alzheimer's disease. Acta Neuropathol 120:185-193. 10.1007/s00401-010-0690-1

Rammensee HG. 1995. Chemistry of peptides associated with MHC class I and class II molecules. Curr Opin Immunol 7:85-96.

Rondini S, Micoli F, Lanzilao L, Hale C, Saul AJ, and Martin LB. 2011. Evaluation of the immunogenicity and biological activity of the Citrobacter freundii Vi-CRM197 conjugate as a vaccine for Salmonella enterica serovar Typhi. Clin Vaccine Immunol 18:460-468. 10.1128/CVI.00387-10

Russo C, Violani E, Salis S, Venezia V, Dolcini V, Damonte G, Benatti U, D'Arrigo C, Patrone E, Carlo P, and Schettini G. 2002. Pyroglutamate-modified amyloid beta-peptides--AbetaN3(pE)--strongly affect cultured neuron and astrocyte survival. J Neurochem 82:1480-1489.

Saido TC, Iwatsubo T, Mann DM, Shimada H, Ihara Y, and Kawashima S. 1995. Dominant and differential deposition of distinct beta-amyloid peptide species, A beta N3(pE), in senile plaques. Neuron 14:457-466.

Schilling S, Lauber T, Schaupp M, Manhart S, Scheel E, Böhm G, and Demuth HU. 2006. On the seeding and oligomerization of pGlu-amyloid peptides (in vitro). Biochemistry 45:12393-12399. 10.1021/bi0612667

Schilling S, Zeitschel U, Hoffmann T, Heiser U, Francke M, Kehlen A, Holzer M, Hutter-Paier B, Prokesch M, Windisch M, Jagla W, Schlenzig D, Lindner C, Rudolph T, Reuter G, Cynis H, Montag D, Demuth HU, and Rossner S. 2008. Glutaminyl cyclase inhibition attenuates pyroglutamate Abeta and Alzheimer's disease-like pathology. Nat Med 14:1106-1111. 10.1038/nm.1872

Schlenzig D, Manhart S, Cinar Y, Kleinschmidt M, Hause G, Willbold D, Funke SA, Schilling S, and Demuth HU. 2009. Pyroglutamate formation influences solubility and amyloidogenicity of amyloid peptides. Biochemistry 48:7072-7078. 10.1021/bi900818a

Schlenzig D, Rönicke R, Cynis H, Ludwig HH, Scheel E, Reymann K, Saido T, Hause G, Schilling S, and Demuth HU. 2012. N-Terminal pyroglutamate formation of Aβ38 and Aβ40 enforces oligomer formation and potency to disrupt hippocampal long-term potentiation. J Neurochem1 121:774-784. 10.1111/j.1471-4159.2012.07707.x

Selkoe DJ. 2001. Alzheimer's disease: genes, proteins, and therapy. Physiol Rev 81:741-766.

Sevalle J, Amoyel A, Robert P, Fournié-Zaluski MC, Roques B, and Checler F. 2009. Aminopeptidase A contributes to the N-terminal truncation of amyloid beta-peptide. J Neurochem 109:248-256. 10.1111/j.1471-4159.2009.05950.x

Shinefield HR. 2010. Overview of the development and current use of CRM(197) conjugate vaccines for pediatric use. Vaccine 28:4335-4339. 10.1016/j.vaccine.2010.04.072

Sullivan CP, Berg EA, Elliott-Bryant R, Fishman JB, McKee AC, Morin PJ, Shia MA, and Fine RE. 2011. Pyroglutamate-Aβ 3 and 11 colocalize in amyloid plaques in Alzheimer's disease cerebral cortex with pyroglutamate-Aβ 11 forming the central core. Neurosci Lett 505:109-112. 10.1016/j.neulet.2011.09.071

Tabira T. 2010. Immunization therapy for Alzheimer disease: a comprehensive review of active immunization strategies. Tohoku J Exp Med 220:95-106.

Vingtdeux V, Chandakkar P, Zhao H, Blanc L, Ruiz S, and Marambaud P. 2015. CALHM1 ion channel elicits amyloid-β clearance by insulin-degrading enzyme in cell lines and in vivo in the mouse brain. J Cell Sci 128:2330-2338. 10.1242/jcs.167270

Vingtdeux V, Davies P, Dickson DW, and Marambaud P. 2011. AMPK is abnormally activated in tangle- and pre-tangle-bearing neurons in Alzheimer's disease and other tauopathies. Acta Neuropathol 121:337-349. 10.1007/s00401-010-0759-x

Winblad B, Graf A, Riviere ME, Andreasen N, and Ryan JM. 2014. Active immunotherapy options for Alzheimer's disease. Alzheimers Res Ther 6:7. 10.1186/alzrt237

Wirths O, Erck C, Martens H, Harmeier A, Geumann C, Jawhar S, Kumar S, Multhaup G, Walter J, Ingelsson M, Degerman-Gunnarsson M, Kalimo H, Huitinga I, Lannfelt L, and Bayer TA. 2010. Identification of low molecular weight pyroglutamate A{beta} oligomers in Alzheimer disease: a novel tool for therapy and diagnosis. J Biol Chem 285:41517-41524. 10.1074/jbc.M110.178707

Wisniewski T, and Goñi F. 2015. Immunotherapeutic approaches for Alzheimer's disease. Neuron 85:1162-1176. 10.1016/j.neuron.2014.12.064

